# Plasmid transfer is biased towards close kin in bacteria from natural populations

**DOI:** 10.1101/434282

**Authors:** Tatiana Dimitriu, Lauren Marchant, Angus Buckling, Ben Raymond

**Affiliations:** Department of Biosciences, University of Exeter, Penryn Campus, Cornwall TR10 9FE, UK

## Abstract

Plasmids play a key role in microbial ecology and evolution, yet the determinants of plasmid transfer rates are poorly understood. Here we investigate the importance of genetic similarity between naturally co-occurring *Escherichia coli* isolates in the transfer of two plasmids (narrow-host-range R1 and broad-host-range RP4). We uncovered extensive variability, spanning over five orders of magnitude, in the ability of isolates to donate and receive plasmids. Overall, transfer was strongly biased towards clone-mates, but not correlated to genetic distance between donors and recipients. Transfer was limited by the presence of a functional restriction-modification system in recipients, thus bias towards kin might be explained by sharing of identical restriction systems. Such conjugation within lineages sets the stage for longer-term pair-wise coevolutionary interactions between plasmids and bacterial hosts.

## Introduction

Conjugative plasmids play a central role in horizontal gene transfer, impacting both evolutionary and ecological processes. At large phylogenetic scales, they are the main vector of genetic exchange among bacteria (Halary et al 2009), shaping gene flow and long-term adaptation of communities. They also encode a diversity of “accessory genes” (Frost et al 2005) often conferring environment-specific adaptations such as antibiotic and metal resistance and virulence traits. As a consequence, the dynamics of plasmid transfer has crucial consequences on the outcome of competition between strains or species, which in turn can both drive the epidemiology of bacterial pathogens and influence ecosystem services. In particular, antibiotic-resistance-conferring plasmid transfer can govern success of strains within patients (Conlan et al 2016, Porse et al 2017, Sheppard et al 2016) and facilitate pathogen epidemics (Baker et al 2018). An understanding of factors controlling transfer rates is therefore critical.

A striking feature of plasmid transmission is its variability. Indeed, estimates of transfer rates draw fundamentally different conclusions about whether plasmids can naturally persist in the absence of selection on plasmid-carried traits (Simonsen et al 1991, Bergstrom et al, 2000, Slater et al 2008), although recent work suggest transfer rates are typically high enough (Lopatkin et al, 2017). Transfer rates are dependent on both the initiation of conjugation in donor cells, and successful establishment in recipient cells (Thomas and Nielsen, 2005). Quantification of transfer rates in the laboratory has mostly focused on a few laboratory strains, despite the strong effect host genotype and plasmid-host interactions can have on plasmid transfer rates. The few data available suggest that laboratory strains have higher than average transfer abilities (Gordon, 1992) making them poor models for natural populations (Hobman et al, 2007). Transfer of plasmid R1 is highly variable among natural isolates, with rates spanning seven orders of magnitude even within species (Gordon, 1992, Dionisio et al, 2002). Host community composition might thus shift plasmid outcome between the two extremes of loss or fixation – with rare efficient donors having a particularly strong effect (Dionisio et al, 2002).

Both donor and recipient abilities might vary independently, but genetic interactions between donor and recipient genotypes will also occur. Particularly, high transfer rates with successful establishment in recipient cells might be favoured by genetic similarity between donors and recipients. Parasite success in new hosts has been correlated to genetic distance between hosts in various organisms, for instance phages (Scanlan et al, 2017) or insect viruses (Longdon et al, 2011). Transfer restriction could contribute to global patterns in plasmid host range, where different plasmid groups are specifically associated to different host lineages (Williams et al 2013, Medaney et al 2016, Méric et al 2018), a pattern also observed for pathogenic lineages (McCarthy and Lindsay 2012, Mathers et al 2015). Transfer to kin i.e. plasmid donors and recipients belonging to the same genotype, was shown to be particularly efficient across a range of hosts from different strain collections (Dimitriu et al 2016). Biased transfer towards kin can promote higher investment in transfer for beneficial plasmids, which could lead to higher transfer for traits including antibiotic resistance (Dimitriu et al, 2016). However, this effect relies on the observed bias in transfer actually occurring among strains coexisting in the field.

To understand the ecological consequences of diversity and biases in transfer, variation needs to be characterized within natural populations and not among strain collections from lineages that are unlikely to coexist in the field. Variation might exist only at a large phylogenetic scale or be the product of environment-specific selective forces, such that strains isolated from the same environment have much more homogeneous transfer rates or display no bias in transfer towards kin. Here, we investigate the transfer rates of two resistance plasmids, R1 and RP4 with respectively narrow and broad host ranges, among a collection of *E. coli* host isolates from cattle, for which population structure and native plasmid content have been characterized previously (Medaney et al, 2016). We ask if the striking diversity in transfer previously observed within laboratory strain collections is still present within populations, when host bacteria are isolated from the same or similar environments and at short phylogenetic scales; and explore genetic factors that could contribute to biased transfer towards kin.

## Methods

### Bacterial strains

*E. coli* field isolates (Table S1) were selected from the field collection characterized in [Medaney et al, 2016], which surveyed *E. coli* diversity and plasmid content in grazing cattle. We selected 14 strains belonging to 7 different serotypes (H-antigen genotype, Wang et al 2003) to cover *E. coli* diversity present in the collection. Isolates with detected plasmid replicons were excluded. To test for both genotype and site effect, we selected strains such that couples of strains sharing the same serotype were isolated from different sites; and reciprocally strains isolated from the same site had different serotypes. We also used the standard laboratory strain *E. coli* K12 MG1655, its mutant MG1655 ∆hsdS::Kn generated by P1 transduction from the Keio collection (Baba et al, 2005) and another MG1655 derivative MFDpir (Babic et al, 2008).

### Plasmids and conjugation assays

We characterized transfer rates for two plasmids belonging to the incompatibility groups most abundant in the field collection (Medaney et al, 2016). R1 plasmid (Meynell and Datta, 1966) is a narrow-host range IncFII plasmid, in which regulation of transfer is representative of the majority of IncF plasmids (Fernandez-Lopez et al 2016). RP4 plasmid (Pansegrau et al, 1994) is a broad-host range IncP plasmid. R1 and RP4 were conjugated into unmarked field isolates using the standard donor strain MFDpir (Babic et al 2008) that requires di-aminopimelate (DAP, Sigma-Aldrich) to grow in LB medium. For all other conjugation assays, spontaneous rifampicin-resistant (Rif^R^) mutants of MG1655 and the 14 field isolates were generated by plating overnight cultures on LB-agar with rifampicin (Rif, Sigma-Aldrich) at 100 µg/mL, to use as recipients. Plasmid-containing bacteria were selected with kanamycin (Kn, 50 ug/mL), except in experiments including MG1655 ΔhsdS::Kan strain and R1 plasmid, for which chloramphenicol (Cm, Sigma-Aldrich, 25 ug/mL) was used instead.

Conjugation assays were performed by mixing equal volumes of overnight cultures of donors and recipients with 10-fold total dilution into 1 mL LB medium, supplemented with DAP 0.3 mM when MFDpir was used as a donor. Overnight cultures for conjugation experiments did not contain antibiotics. Mixes were incubated at 37°C with 150 rpm shaking. To favor detection of relatively low transfer rates, mating assays were performed for 3h as a general standard. When stated, assays with a reduced 1h mating were performed in order to limit secondary transfer from transconjugants. Donor, recipient and transconjugant densities were then estimated by dilution plating onto respectively LB-agar (LBA)+DAP+Kn, LBA and LBA+Kn (assays with MFDpir as a donor) or LBA+Kn/Cm, LBA+Rif and LBA+Rif+Kn/Cm (other assays). Each conjugation was performed with at least two replicates per experiment and two independent experiments. Control assays with donors only and recipients only were done in parallel, and never showed any growth on selective transconjugant plates.

### Isolate genotyping and phylogenetic distance

We used the set of three phylogenetic markers identified in (Sahl et al, 2012) to study phylogenetic relationships among *E. coli* isolates. The 3 markers *dinG*, *DPP* and *tonB* were amplified from the 14 field isolates with PCRBio Taq Mix Red (PCR Biosystems) using primers described in Sahl et al, (2012), and sequenced through Eurofins Genomics. For the laboratory strains MG1655 and its derivative MFDpir, sequences from GenBank accession NC_000913 were used. Sequence data were pre-processed in Geneious (version 8.1.6). Amplicons were trimmed at both the 3’ and 5’ ends, to remove low quality sequence i.e. base pairs with an error probability above 5%. High quality alignments (respectively 854, 792 and 725 bp long for *dinG*, *DPP* and *tonB*) were concatenated and used to determine multi-locus phylogenetic distance. For isolates D7.8 and oc5.1, *tonB* primers did not yield any product; *DPP* and *dinG* products revealed both isolates were part of *E. marmotae* species. Phylogenetic trees were made with a weighted neighbor-joining tree building algorithm implemented in Geneious. To obtain phylogenetic distances among all isolates including *E. marmotae*, the tree build with *DPP* and *dinG* products only was used; distances obtained were highly correlated with the ones using all three gene products within *E. coli* isolates (Spearman correlation coefficient ρ = 0.98, p<2.2 10^−16^).

### Data analysis

Conjugation rates were measured as 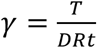 (mL.cell^−1^.h^−1^) where T, D, and R respectively indicate the density of transconjugants, donors and recipients (cells.mL^−1^), and t indicates incubation time (h). When no transconjugants were detected, a threshold conjugation rate was calculated by assuming that one single transconjugant colony was observed. Note that this threshold rate will vary across biological replicates because of variation in donor and recipient cell counts. For assays using field isolates as both donors and recipients, variable growth was observed and data were excluded when either recipient or donor densities were less than 2 10^7^ cells.mL^−1^, in order to limit variation in computed conjugation rates due to low donor or recipient densities. As transfer rate values spanned several orders of magnitude, all statistical analysis was done on log10-transformed data. R Version 3.4.1 was used for all analyses (R Development Core Team; http://www.r-project.org).

## Results

### Variation in donor and recipient ability across natural isolates

To analyze the amplitude of variation in transfer rates, we first measured transfer rates using K-12 laboratory strain MG1655 derivatives as standard donor or recipient. Conjugation assays were performed with each 14 field isolates as donor to, or recipient from K-12. One of 14 isolates, D2.2 was observed to repeatedly kill K-12 strains in each assay (with less than 1% of K-12 inoculum detected after mating), it was thus excluded from analysis.

For both R1 and RP4 plasmids, transfer rates spanned more than 5 orders of magnitude overall, from around 10^−16^-10^−15^ mL/cell/h (detection threshold) to more than 10^−10^ mL/cell/h (Figure 1). For the isolates tm8.6 and R1.9, we were unable to obtain any clones with RP4 across several assays, revealing very low recipient ability; donor ability was not quantified. For both plasmids, donor and recipient ability of K-12 was always as high or higher than any of the field isolates. Excluding K-12, recipient genotype significantly affected conjugation rate from a standard donor for both R1 (*F*_12,39_ = 16.05, *p* < 2.10^−11^) and RP4 (*F*_12,39_ = 13.1, *p* < 10^−9^). Similarly, excluding K-12 donor genotype significantly affected conjugation rate towards the standard recipient for both R1 (*F*_12,37_ = 11.3, *p* < 10^−8^) and RP4 (*F*_10,33_ = 8.07, *p* = 2.10^−6^). The lowest amplitude of variation was observed for R1 plasmid donor ability (Figure 1), for which all measured rates were above 10^−14^ mL/cell/h. We hypothesized that efficient secondary transfer from MG1655 recipients was masking actual variability in transfer from primary donors. Measuring conjugation rates with reduced mating time revealed higher variation among isolates (Figure S1) and a stronger effect of the donor (*F*_12,37_ = 31.8, *p* < 10^−15^).

**Figure 1:**
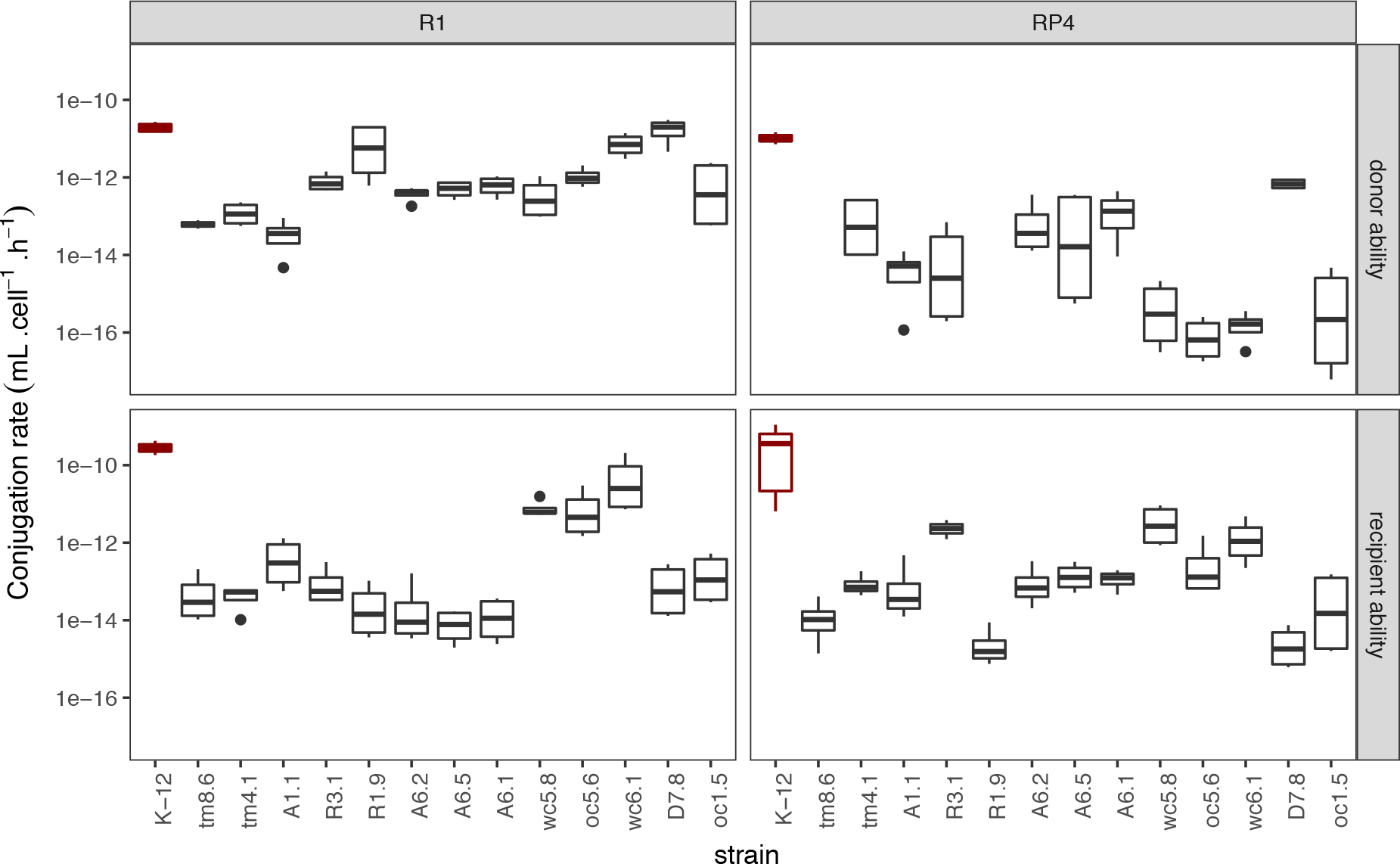
Extensive variation in plasmid donor and recipient ability across field isolates of *E. coli*. Conjugation rates were measured in liquid with shaking over 3h, towards K-12 MG1655 Rif^R^ (donor ability, top) and from K-12 MFDpir (recipient ability, bottom). Rates with K-12 as both donor and recipient are shown on the left (red); field isolates are then ordered by their phylogenetic distance to K-12. Isolates which names begin with the same letters indicate shared site of isolation.

No correlation was observed among isolates between average donor and recipient ability (Spearman rank-correlation ρ = 0.02, *p* = 0.92); or between average rates between plasmids R1 and RP4 (Spearman rank-correlation ρ= 0.35, *p* = 0.08). Moreover, phylogenetic distance from the standard K-12 donor or recipient did not explain average conjugation rates towards or from natural isolates (Figure S2, R^2^ = 0.003, *F*_1,48_ = 0.14, *p* = 0.71).

### Transfer within and between kin

To explore diversity in transfer rates among field isolates and test if transfer is increased towards kin, we performed for each donor conjugation assays to a marked recipient of the same isolate (transfer to kin); and to a randomly chosen isolate both belonging to a different serotype and isolated from a different field site (transfer to non-kin). Rates of transfer were strongly variable across isolates, spanning 5 orders of magnitude from < 10^−16^ (detection threshold) to > 10^−12^ mL. cell^−1^.h^−1^ (Figure 2).

**Figure 2:**
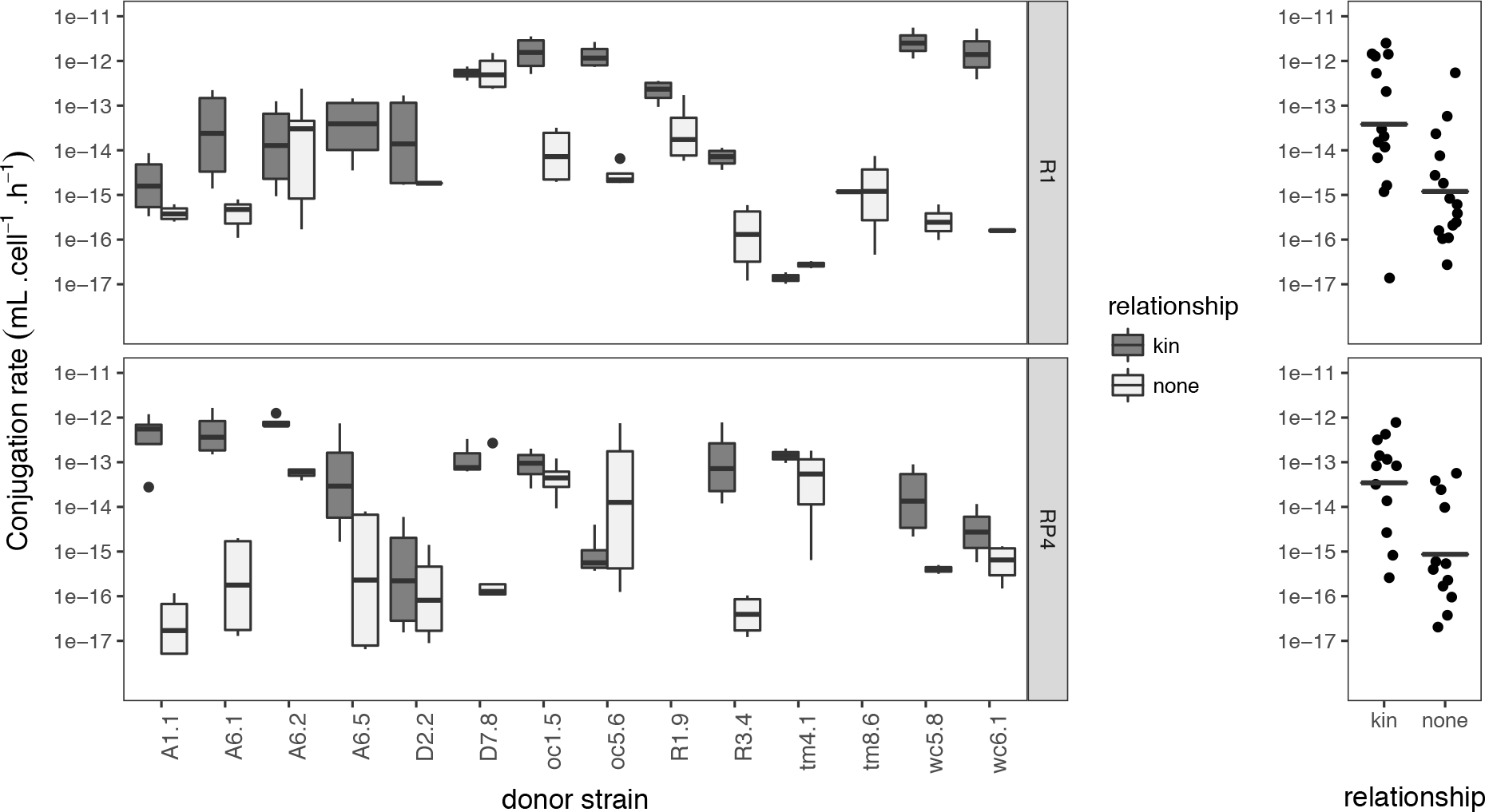
Transfer rates among field isolates and bias towards kin. Mating assays were performed for 3h, with donors containing either R1 or RP4 plasmid, and shown ordered by donor isolate, with the recipient isolate being the same as the donor (kin, black) or another field isolate (non-kin, light gray, see Table S1 for recipient identity). Summary graphs at the right show average transfer per couple of strains (dots) and overall geometric means per treatment (lines).

For each plasmid, the effect of relationship between isolates (kin or non-kin) was the main effect controlling conjugation rates: for R1, *F*_1,70_ = 108, *p* < 10^−15^; for RP4, *F*_1, 74_ = 34.1, *p* < 2 10^−7^. In addition, the identity of donor and recipient also had a strong effect (for R1 plasmid, donor effect *F*_13,70_ = 20, *p* < 2.10^−16^, recipient effect *F*_9,70_ = 10.9, *p* < 3.10^−10^; for RP4 plasmid, donor effect *F*_11,74_ = 5.93, *p* < 10^−6^, recipient effect *F*_7,74_ = 11.5, *p* < 10^−9^). On average, a given donor transferred plasmid R1 towards kin 28.7-fold more efficiently than towards non-kin, and plasmid RP4 39.9-fold more efficiently. This effect was highly variable across isolates; however, no isolate was observed to transfer at significantly higher rates towards non-kin. There was still high variability among isolates considering only transfer towards clone-mates (for R1, *F*_13,35_ = 14.1, *p* = 3.10^−10^, for RP4 *F*_11,33_ = 9.09, *p* < 10^−6^), still spanning 5 orders of magnitude. When the same couples of isolates were tested for both plasmids, no correlation in average transfer rates between R1 and RP4 was observed across couples (Pearson correlation coefficient r_19_ = 0.17, *p* = 0.46).

### Transfer within and between serotypes and sites

To investigate in more detail what governs the higher conjugation rates observed among clone-mates, we performed additional conjugation assays, focusing on R1 plasmid. We chose couples of isolates that share serotype (initial assessment of their relatedness) or field site of isolation (Table S1). We asked if similar variation is observed with shared serotype or within sites, or if isolates from the same field or with same serotype transfer preferentially to each other. Overall, the effect of relationship between donors and recipients (clone-mates, no relation, same serotype or same field) was still strong (transfer rate ~ donor strain + recipient strain + relationship, *F*_3,142_ = 44.95, *p* < 2.10^−16^). However, post-hoc Tukey tests showed that the only relationship that was significantly different from others was clone-mates, with higher transfer rates (*p* < 10^−7^ for all comparisons). Isolates sharing serotype or site of isolation or none were similar (*p* > 0.5 for all comparisons). Donor and recipient identity also had a strong effect (respectively *F*_13,142_ = 14.2, *p* < 2.10^−16^, and *F*_13,142_ = 10.2, *p* < 10^−14^), and isolates from the same field sites also showed variation in transfer rates (Figure S3).

Transfer rates within serotypes were not higher than among non-related isolates, suggesting that sharing serotype does not confer high enough relatedness to be equivalent to kin. That could be due to serotypes encompassing large genetic distances, or to kin discrimination requiring extremely close relatedness. To distinguish between those cases, we derived phylogenetic distance among isolates, which revealed that serotype was a poor marker of phylogenetic distance (Figure 3B). Two isolates D7.8 and oc5.1 were even identified as belonging to *E. marmotae*, despite sharing serotypes with *E. coli* isolates. On the other hand, three closely related isolates from the same cow pat (A6.1, A6.2 and A6.5) belonged to different serotypes. Overall, there was a small but significant negative effect of phylogenetic distance on conjugation rates (estimate = −9.03±2.86, r^2^=0.05, *p* < 0.002). However, when including kin / non-kin status in the model, the effect of phylogenetic distance disappeared (transfer rate ~ kin status + distance, distance estimate 0.65±3.14, *p* = 0.836). Transfer rates are thus not linked to general phylogenetic similarity, but only couples with strict kin identity have overall statistically higher transfer rates. Among non-kin, both closely related and inter-species couples had transfer rates spanning from <10^−16^ to > 10^−13^ mL/cell/h (Fig 3C).

**Figure 3:**
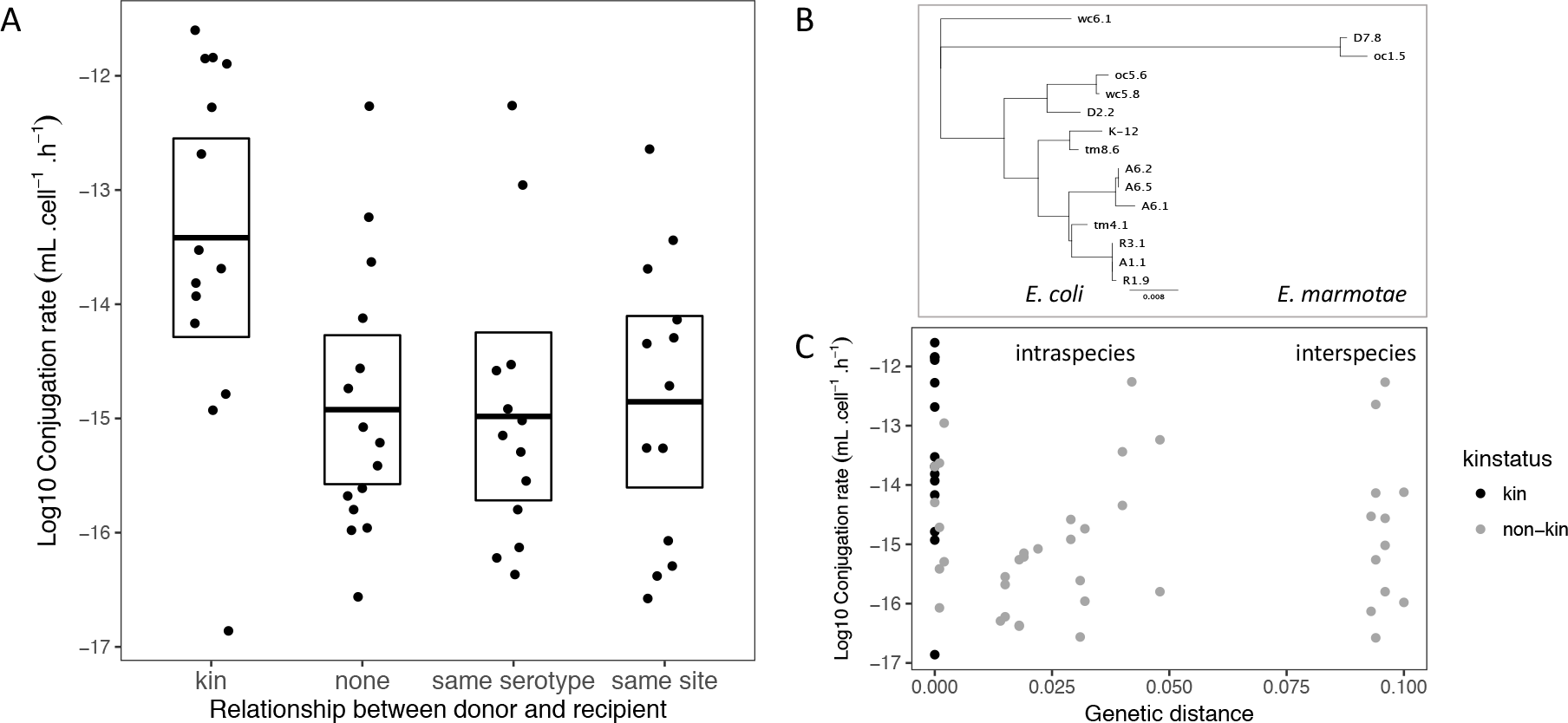
Higher transfer rates among field isolates are not correlated to phylogenetic distance but strictly restricted to kin (clone-mates) The average R1 plasmid conjugation rate for a given couple of donor and recipient isolates is shown function of the relationship initially defined between donor and recipient (A), and of the genetic distance between donor and recipients (C). Phylogenetic relationships between 14 field isolates and K-12 laboratory strain are shown in B.

### Variation in restriction-modification systems as a mechanism for biased transfer

The restriction to clone-mates of high transfer rates suggests that barriers to transfer are not simply a function of genetic similarity. One barrier could be variation in restriction-modification (RM) systems, which can lead to decrease in transfer rates (Roer et al., 2015): RM systems are based on the combination of a restriction enzyme cleaving a specific DNA sequence, and a cognate methyl-transferase protecting that same sequence. Foreign DNA originating from cells without the same RM system is thus targeted by restriction. To test if restriction-modification plays a role in transfer variation, we compared transfer rates of the field isolates towards the standard K-12 recipient, MG1655 (RM^+^) and its mutant with no functional type I RM system (RM^−^, Sain and Murray, 1980), MG1655 ∆hsdS::Kan. We hypothesized that transfer would be overall higher towards the RM^−^ strain, if restriction efficiently targets R1 and enough field isolates do not possess a system with same specificity.

We first confirmed that R1 transfer within K-12 is affected by restriction (Figure 4, left): as expected, the RM^+^ strain transfers equally well to both RM^+^ and RM^−^ recipients; the RM^−^ strains transfers plasmids at the same rates towards itself, but transfer from the RM^−^ strain is restricted in RM^+^ recipients. When measuring transfer from field isolates (Figure 4, right), in addition to a strong effect of donor isolate (*F*_12,106_ = 47.2, *p* < 2 10^−16^), recipient RM status was also significant (*F*_1,106_ = 30.6, *p* < 3 10^−7^). On average, the RM^−^ recipient received R1 plasmid at 3.15-fold higher rates than the RM^+^ recipient. However, the donor/recipient interaction was significant as well (*F*_12,106_ = 2.75, *p* = 0.003), with only some isolates transferring R1 more efficiently towards the RM^−^ strain, as expected if some donors also bear a RM system with same specificity as K-12.

**Figure 4:**
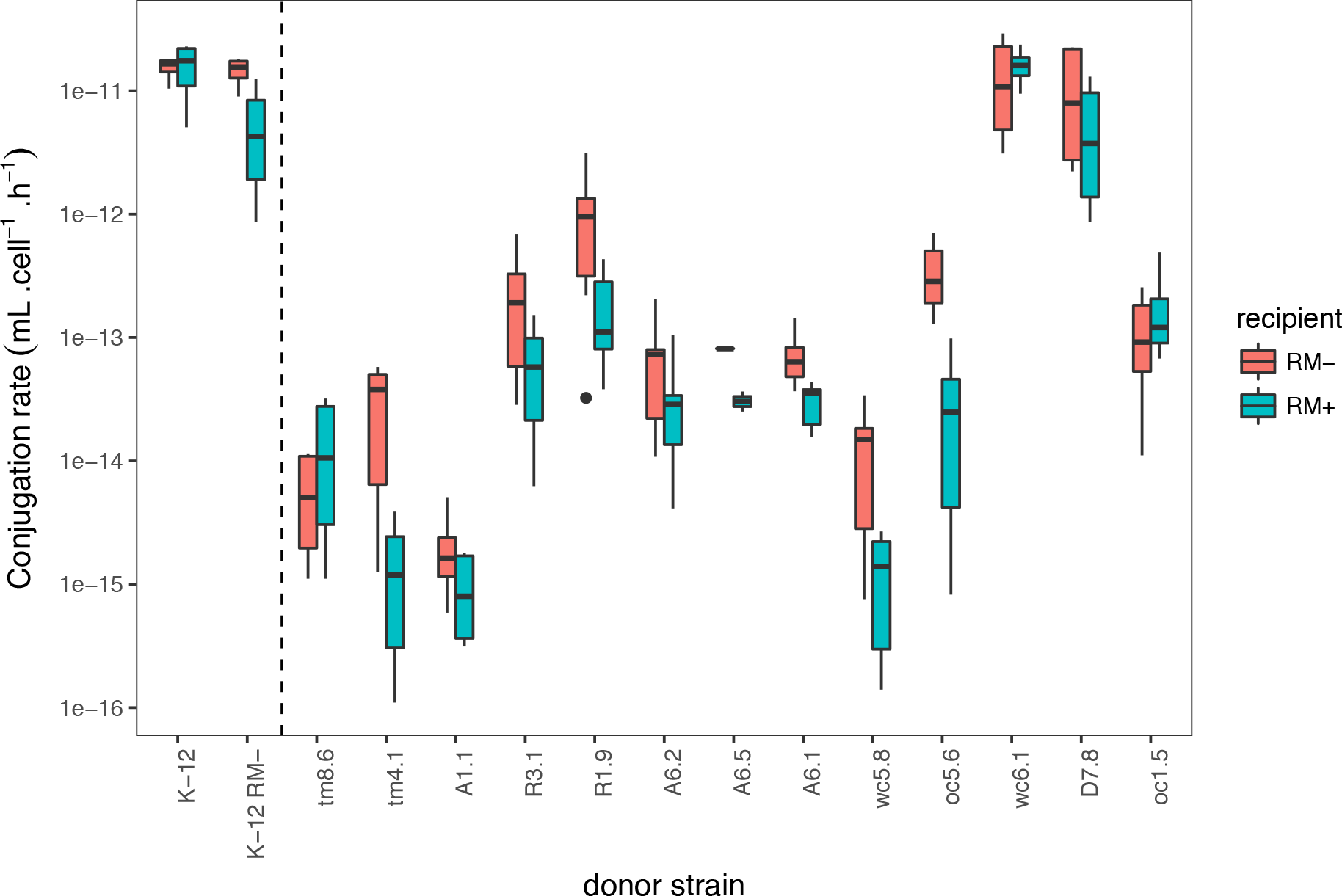
Effect of the EcoKI restriction-modification system on transfer from natural isolates. Mating assays were performed for 1h, from R1 plasmid donors shown on the x-axis towards a K-12 recipient with (RM^+^, blue) or without (RM^−^, red) its native RM system EcoKI. Positive controls are shown on the left: deactivating RM in donors decreases conjugation rate when recipients are RM-positive.

## Discussion

Large variation in plasmid transfer rates has already been described for R1 plasmid within *E coli* (Gordon 1992, Dionisio et al, 2005). We show here that the same variation still occurred within a set of isolates from common environments, implying that it does not only arise from different environment-dependent selective pressures. We also observed cases of large differences in transfer rate within field site or for closely related isolates. Donor and recipient abilities were not correlated, consistent with different mechanistic basis. For recipient ability, we identified restriction-modification as a likely mechanism, and other defence systems, for instance CRISPR-Cas may be involved too (Thomas and Nielsen, 2005; Price et al 2016). Transfer rates for R1 and RP4 plasmids were not correlated either, consistent with a different regulation of transfer operons (Frost and Koraimann, 2010). Interestingly, broad-host range RP4 had similar variation in transfer among hosts, and was no less sensitive to host control than R1, despite suggestions that IncF narrow host range might arise from their more complex regulation by host cells (de la Cruz et al, 2014). Finally, variation in transfer rates among natural isolates might even be higher than estimated here, as we selected isolates with no detected replicons, limiting the effect of modulation of transfer rates by co-resident plasmids (Gama et al, 2017).

The laboratory K-12 strain, MG1655, had transfer rates much larger than the field isolate average for both plasmids. High recipient ability is in agreement with a previous study that found very few genes in K-12 able to limit recipient ability (Pérez-Mendoza et al, 2009). MG1655 has an active restriction system, and recipient ability is increased in the *hsdS* gene mutant, but natural isolates likely possess more active defence mechanisms. Similar results were obtained with phages, where MG1655 was found to be susceptible to many more phages than natural *E. coli* isolates (Allen et al, 2016). Interactions between plasmids and other elements like prophages might also affect transfer operon expression (Miller et al, 1985) or plasmid cost (San Millan et al, 2015) and explain why donor ability is lower than in K-12 for the majority of field isolates, that probably bear an abundance of prophages and other mobile elements.

In addition to donor and recipient identity, the main factor controlling transfer rates was the nature of the genetic interaction between them, with transfer towards kin (clone-mates) being more than 10-fold higher than towards non-kin. We therefore extend the pattern identified previously (Dimitriu et al, 2016) to a second plasmid, the broad-host-range RP4. The average bias towards kin was even higher for RP4, consistent with the fact that it lacks anti-restriction genes present on R1 (Chilley and Wilkins, 1995). Importantly, we show that bias towards kin occurs among *E. coli* lineages co-existing in the field, sometimes isolated from the same cow pat, indicating that this phenomenon is prevalent in natural populations. Moreover, we demonstrate here that the effect is restricted to close kin, i.e. isolates from the same genotype, and transfer is not higher towards isolates with relatively closer genetic distance, or isolated from the same site. Thus, discrimination towards kin is here not a function of average genetic distance among strains (Ostrowski et al, 2008), but might arise from a combination of few loci (Lyons et al, 2016).

We identified restriction-modification as a likely mechanism contributing to discrimination in transfer, showing that recipients with inactivated RM receive R1 plasmid at higher rates from field isolates. Restriction was previously shown to limit plasmid conjugation rates with relatively low efficiency (Roer et al, 2015), likely because the first transconjugants that escaped restriction are then protected from further restriction when transferring to kin. RM systems are tremendously variable and mobile among *E. coli* isolates (Murray, 2002; Barcus et al, 1995). Expression of several RM systems can have a multiplicative effect on restriction efficiency (Schouler et al, 1998), thus the effect we measure with a single RM system could be amplified among diverse isolates. Our results are in agreement with studies describing the role of RM systems in restricting transfer among lineages in several species (Waldron and Lindsay, 2006; Oliveira et al, 2016). RM variation is the main known mechanism that could lead to specific discrimination in conjugative process. However, other discrimination or structuring processes, not directly targeting plasmid conjugation, would also lead to discrimination in transfer if they affect how much donors encounter kin vs. non-kin. This includes non-kin killing by bacteriocins, a form of kin discrimination (Strassmann et al, 2011). Spatial structure, that promotes transfer to kin in the absence of discrimination mechanisms (Dimitriu et al, 2016), can also bias transfer across a population. Indeed the *E. coli* populations sampled for this study show strong population structure, indicating that opportunities for transfer to plasmid null isolates occur predominantly within genotypes (Medaney et al 2016).

The diversity in transfer rates that we uncover has consequences for understanding plasmid maintenance and ecological dynamics. The rates of transfer within each lineage, one of the key determinants of plasmid maintenance (Stewart and Levin, 1977), vary here by five orders of magnitude. Plasmids from 9 different groups were recently shown to be transferred at rates sufficient for persistence, in a classical K-12 strain (Lopatkin et al, 2017). Our results suggest that these conclusions should be taken with caution, as natural *E. coli* will likely transfer less than K-12. The scale of variation we observe implies that maintenance of plasmids might depend on subtle details of host community composition. Still, a few isolates transfer plasmids almost as efficiently as K-12, and efficient donors can promote transfer in mixed bacterial populations (Dionisio et al, 2002), helping maintaining plasmids in mixed communities (Hall et al, 2016). On the other hand, the biased transfer to kin we observe will limit that dynamics, and promote plasmid clustering in distinct lineages. Moreover, when plasmids confer benefits to their hosts, as with antibiotic exposure for antibiotic resistance plasmids, restricting transfer towards kin will be beneficial to host bacteria. It will also promote indirect selection of efficient donor hosts, through kin selection mechanisms (Dimitriu et al, 2016). However, transfer towards non-kin, that is efficient for some pairs of isolates in our field collection, might also benefit the hosts when the transferred plasmids bear public-good encoding genes (Dimitriu et al, 2014, 2018), for instance virulence, antibiotic resistance or detoxification genes (Rankin et al, 2010, Raymond et al, 2012). More generally, transfer being the highest within kin, together with the observation that plasmids are not at fixation within serotypes (Medaney et al, 2016) suggests that most plasmid dynamics might actually occur not between lineages (the events most easily detected) but within lineages, with recurring loss or counter-selection of plasmid-bearing cells alternating with selection and plasmid transfer.

## Acknowledgments

We thank Chantal Lotton for P1 transduction, and Andrew Matthews for comments and advice on statistical and sequence analysis.

